# Scaling single cell transcriptomics through split pool barcoding

**DOI:** 10.1101/105163

**Authors:** Alexander B. Rosenberg, Charles M. Roco, Richard A. Muscat, Anna Kuchina, Sumit Mukherjee, Wei Chen, David J. Peeler, Zizhen Yao, Bosiljka Tasic, Drew L. Sellers, Suzie H. Pun, Georg Seelig

## Abstract

Constructing an atlas of cell types in complex organisms will require a collective effort to characterize billions of individual cells. Single cell RNA sequencing (scRNA-seq) has emerged as the main tool for characterizing cellular diversity, but current methods use custom microfluidics or microwells to compartmentalize single cells, limiting scalability and widespread adoption. Here we present Split Pool Ligation-based Transcriptome sequencing (SPLiT-seq), a scRNA-seq method that labels the cellular origin of RNA through combinatorial indexing. SPLiT-seq is compatible with fixed cells, scales exponentially, uses only basic laboratory equipment, and costs one cent per cell. We used this approach to analyze 109,069 single cell transcriptomes from an entire postnatal day 5 mouse brain, providing the first global snapshot at this stage of development. We identified 13 main populations comprising different types of neurons, glia, immune cells, endothelia, as well as types in the blood-brain-barrier. Moreover, we resolve substructure within these clusters corresponding to cells at different stages of development. As sequencing capacity increases, SPLiT-seq will enable profiling of billions of cells in a single experiment.

---

Over three hundred years have passed since the discovery of the cell, yet we still do not have a complete catalogue of cell types or their functions. While transcriptomic profiling of individual cells has emerged as a promising solution to characterizing cellular diversity (1, 2), increases in throughput are needed before a complete “atlas” of cell types can be generated. Recent single cell RNA-seq (scRNA-seq) methods have profiled tens of thousands of individual cells (3–6), revealing new insights about the immune system (7) and identifying new cell types in the brain (8–11). However, since these methods require cell sorters and custom microfluidics or microwells, throughput is still limited, experiments are costly, and access is limited to a small number of labs.

Here we introduce Split Pool Ligation-based Transcriptome sequencing (SPLiT-seq), a low-cost, scRNA-seq method that enables transcriptional profiling of hundreds of thousands of fixed cells in a single experiment using only basic laboratory equipment. In order to process large numbers of cells together, but to retain cell-of-origin information for each transcript, the existing scRNA-seq methods physically separate individual cells into different compartments, in which transcripts are labeled with cell-specific barcodes. Here, we use the cells themselves as containers and label intracellular transcripts using combinatorial indexing (3, 12, 13). In practice, cells are split into different wells, a well-specific barcode is appended to intracellular transcripts, and then cells are pooled back together. By repeating this process several times, we ensure that each cell travels through a unique combination of wells with very high likelihood. Consequently, each transcript will contain a combination of well-specific barcodes indicating its cellular origin. Whereas previous methods scale linearly with the number of available compartments and barcodes, SPLiT-seq scales exponentially with the number of indexing rounds, enabling a rapid scale-up of the number of cells that can be assayed (Fig. 1A) and a reduction in cost per cell to below one cent (Fig. 1B).

**Fig. 1.**
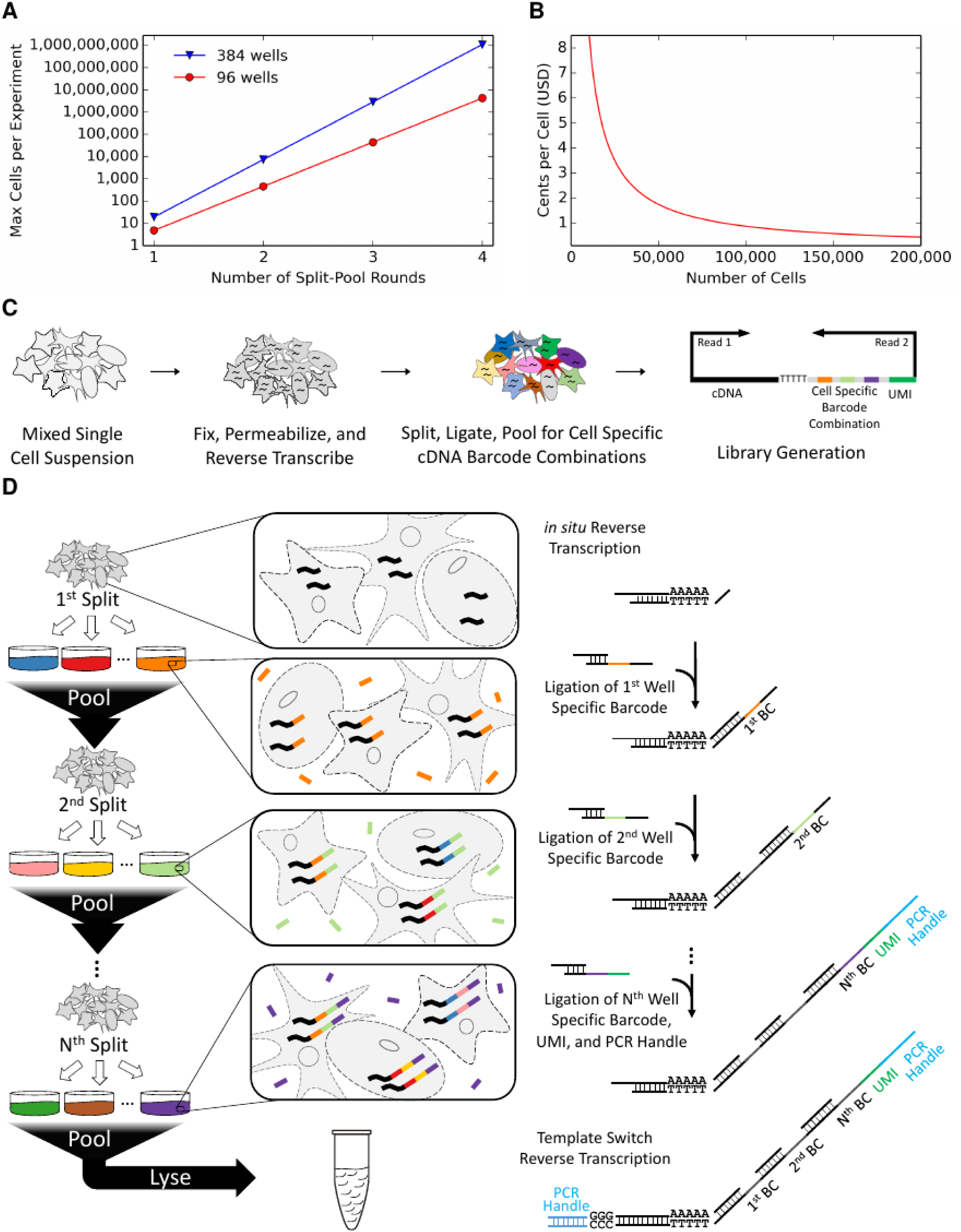
Overview of SPLiT-seq. **(A)** Exponential scalability of SPLiT-seq with number of split-pool rounds. The maximum number of cells is calculated with the assumption that the number of barcode combinations must be twenty times greater than the number of cells. **(B)** Cost per cell for SPLiT-seq. As more cells are processed, costs drop below 1 cent per cell, making SPLiT-seq a cost-effective platform to profile large quantities of cells. **(C)** SPLiT-seq workflow. A mixed single cell suspension is fixed and permeabilized before an *in situ* reverse transcription reaction. Repeated split-ligate-pool rounds then label each cDNA molecule with a cell-specific combination of barcodes. Sequencing libraries are generated from the barcoded cDNA with the first read covering the transcript and the second read covering the UMI and cell-specific barcodes. **(D)** Split-ligate-pooling rounds in SPLiT-seq. After *in situ* reverse transcription of RNA to cDNA, cells are randomly split between wells that each contain a unique well-specific barcode. These barcodes are hybridized and ligated to an overhang on the reverse transcription primer. Cells are pooled back together and randomly split into another set of wells, also containing well-specific barcodes. Second round barcodes are then ligated to the 5’ end of the first round barcode. The cells are pooled back together and subsequent split-ligate-pool rounds can be performed. After *N* rounds, cDNA molecules contain a cell-specific combination of barcodes, a unique molecular identifier, and a universal PCR handle on the 5’ end. A template switching reaction following lysis adds an additional PCR handle on the 3’ side, allowing for cDNA library amplification and generation of sequencing products.

The SPLiT-seq workflow begins with an *in situ* reverse transcription reaction that converts RNA to cDNA within formaldehyde fixed cells in suspension (Fig. 1C). The cells are then subjected to a combinatorial indexing procedure consisting of several split-ligate-pool rounds. To initiate the first round of indexing, cells are randomly distributed into a 96 well plate, with each well containing a unique DNA barcode (Fig. 1D). An *in situ* ligation links the well-specific DNA barcode to the 5’ end of the cDNA molecules. Unligated DNA barcodes are sequestered by subsequent addition of a blocking strand to prevent nonspecific ligations before cells are pooled back together. The split-ligate-pool procedure is then repeated, appending another well-specific barcode to the 5’ end of the first round barcode. After a third and final round of indexing, each cDNA now has 3 barcodes, yielding 884,736 (96^3^) potential barcode combinations. The barcodes in the final round contain unique molecular identifiers (UMI), to ensure that each cDNA molecule is only counted once. Cells with uniquely labelled cDNAs are lysed and can either be stored or immediately converted to expression libraries using a modified existing protocol (4). Importantly, the entire labelling workflow consists just of pipetting steps and no complex instruments are needed.

Cells that have undergone combinatorial indexing can also be aliquoted into multiple sublibraries before lysis. While not necessary, creating sublibraries provides two advantages: First, it makes it possible to sequence a small fraction of cells from a large library before committing to sequencing the entire library. Second, more cells can be sequenced in a single experiment because each sublibrary can be barcoded with an additional PCR index, further increasing the number of combinatorial barcodes.

As a first test of SPLiT-seq’s ability to generate uniquely barcoded cells (UBCs), we performed a species-mixing experiment. We mixed mouse (NIH/3T3) and human (Hela-S3) cells at approximately equal proportions and then used SPLiT-seq to generate a scRNA-seq library with 629 UBCs (Fig. 2A). The library was sequenced and reads were aligned to a combined mouse-human genome. From the alignment, we found that 95.5% of the UBCs were unambiguously generated from a single species (>90% of reads aligned to a single genome) with the remaining 4.5% of cell constituting barcode collisions. However, we expect the true fraction of barcode collisions to be twice the observed fraction, because we cannot identify collisions between two cells from the same species. The species purity in both human and mouse UBCs was high: 98.7% of reads in mouse UBCs and 99.7% of reads in human UBCs aligned to their respective genomes (Fig. 2B). At a relatively shallow sequencing depth of 38,468 reads per cell, we identified a median of 2,382 UMIs and 1,626 genes per human cell and 1,047 UMIs and 762 genes per mouse cell (Fig S1).

**Fig. 2.**
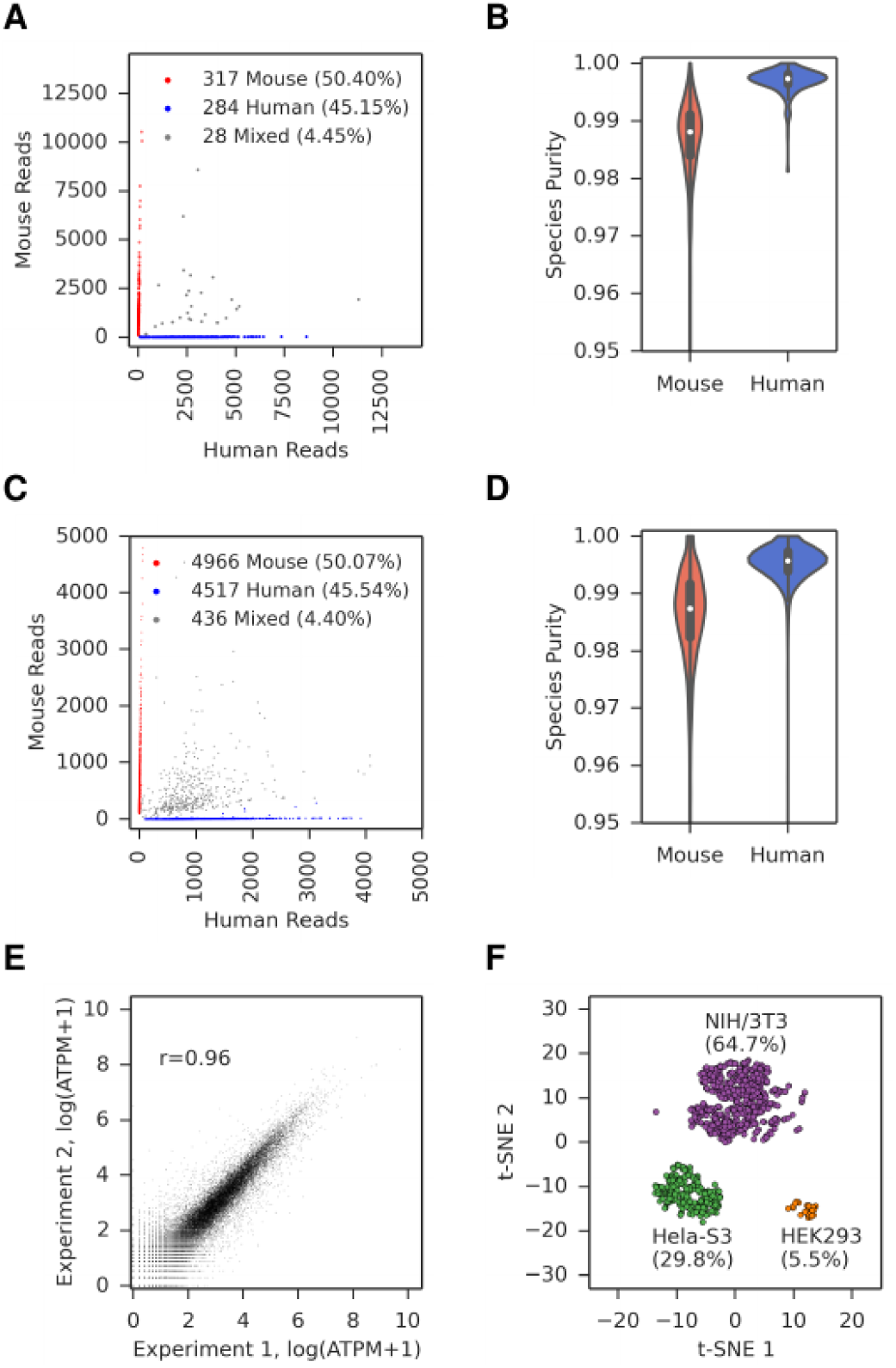
Evaluation of SPLiT-seq. **(A)** The number of human and mouse UMI counts for 629 UBCs. Human UBCs are colored in blue, mouse UBCs are colored in red, and mixed species UBCs are colored in gray. The estimated barcode collision rate is 8.9%. **(B)** Fraction of reads mapping to the correct species for both mouse UBCs and human UBCs. The average purity is 99.7% in human UBCs and 98.7% in mouse UBCs. **(C)** A sublibrary with 9,919 UBCs maintains a similar estimated barcode collision rate (8.8%) as the smaller sublibrary. **(D)** A sublibrary with 9,919 UBCs has a purity of 99.5% in human UBCs and 98.6% in mouse UBCs. **(E)** Biological replicate experiments from different Hela-S3 cells performed multiple weeks apart are highly reproducible (Pearson-r: 0.96). Average gene expression across all cells (log average transcripts per million) are plotted for each experiment. **(F)** SPLiT-seq correctly measures the proportions of subpopulations in a cell mixture. Hela-S3, HEK293, and NIH/3T3 cells were mixed together with input proportions of 27%, 5%, and 67%, respectively. Three clusters were created after running PCA and t-SNE on the gene counts for each UBC. The cluster sizes for each cell line were within 10% of the input proportions.

To evaluate the reproducibility and quantify batch-effects (14) of SPLiT-seq, we repeated the species-mixing experiment several times and generated expression libraries from approximately 10,000 (Fig. 2C) and 25,000 (Fig. S2A) cells. The estimated barcode collision fraction (10k: 8.8%, 25k: 6.6%) and average purity in mouse and human UBCs (10K cells: 98.6%/99.5%, Fig. 2D; 25K: 97.8%/99.1%, Fig. S2B) remained similar to the 629 UBC species-mixing experiment. Gene expression was highly correlated between different sublibraries prepared from the same experiment (Pearson-r: 0.98, Fig. S3) as well as between libraries prepared from different experiments performed several weeks apart (Pearson-r: 0.96, Fig. 2E). In all of our experiments, we observed a substantial enrichment of unspliced, nuclear transcripts (Fig. S4). As other groups have also observed, including these nuclear transcripts improves identification of subpopulations of cells (15).

We then asked whether SPLiT-seq could be used to quantitate proportions of multiple cell types in a heterogeneous population. We first mixed uneven proportions of two human cell lines (5% HEK293, 27% Hela-S3) and a mouse cell line (67% NIH/3T3). Cells were then processed using SPLiT-seq, generating an expression library of 924 UBCs. Principle component analysis (PCA) followed by t-distributed Stochastic Neighbor Embedding (t-SNE) (16) identified three distinct clusters corresponding to HEK293 (5.5 %), Hela-S3 (29.8 %), and NIH/3T3 (64.7 %), all within 10% of the input fractions (Fig. 2F).

For the first application of SPLiT-seq, we chose to profile the developing brain in the early postnatal mouse. In the first week after birth, the mouse brain is rapidly changing; microglia are migrating into the brain, neuroblasts are moving through the rostral migratory stream, and proliferating glial cells are defining the early structure of the brain (17– 18 19). To capture the full view of these transitions, it is necessary to profile the whole brain rather than specific regions or subtypes of cells. However, limits in the scalability of previous scRNA-seq methods have prevented this type of global analysis.

We dissociated an entire brain from a postnatal day 5 mouse and then used SPLiT-seq to generate a scRNA-seq library. After computationally filtering UBCs for potential barcode collisions, we retained 109,069 cell transcriptomes. We performed PCA on the resulting gene counts and mapped the top 16 components into two dimensions using t-SNE. To generate high-level cluster assignments, we fit a Gaussian Mixture Model and then merged clusters based on visual inspection of the t-SNE projection, resulting in 13 distinct clusters (Fig. 3A). We found many genes that were primarily expressed only in a single cluster (Fig. 3B, Fig. S5), of which many corresponded to known cell type markers (20). Over 70% of cells mapped to five different neuronal clusters comprising GABAergic, glutamatergic, granule, Pax2-expressing, and Cntn5-expressing cells. Expression of the proliferation marker *Mki67* revealed that granule cells were still actively dividing whereas cells from all other neuronal clusters had become postmitotic (Fig. S6) (21). The largest non-neuronal clusters corresponded to astrocytes, microglia/macrophages, and oligodendrocytes, which each accounted for 6-7% of all profiled cells. The remaining cells mapped to vascular and leptomeningeal, endothelial, pericyte, ependymal, and Bnc2-expressing clusters, each comprising less than 3% of all cells.

**Fig. 3.**
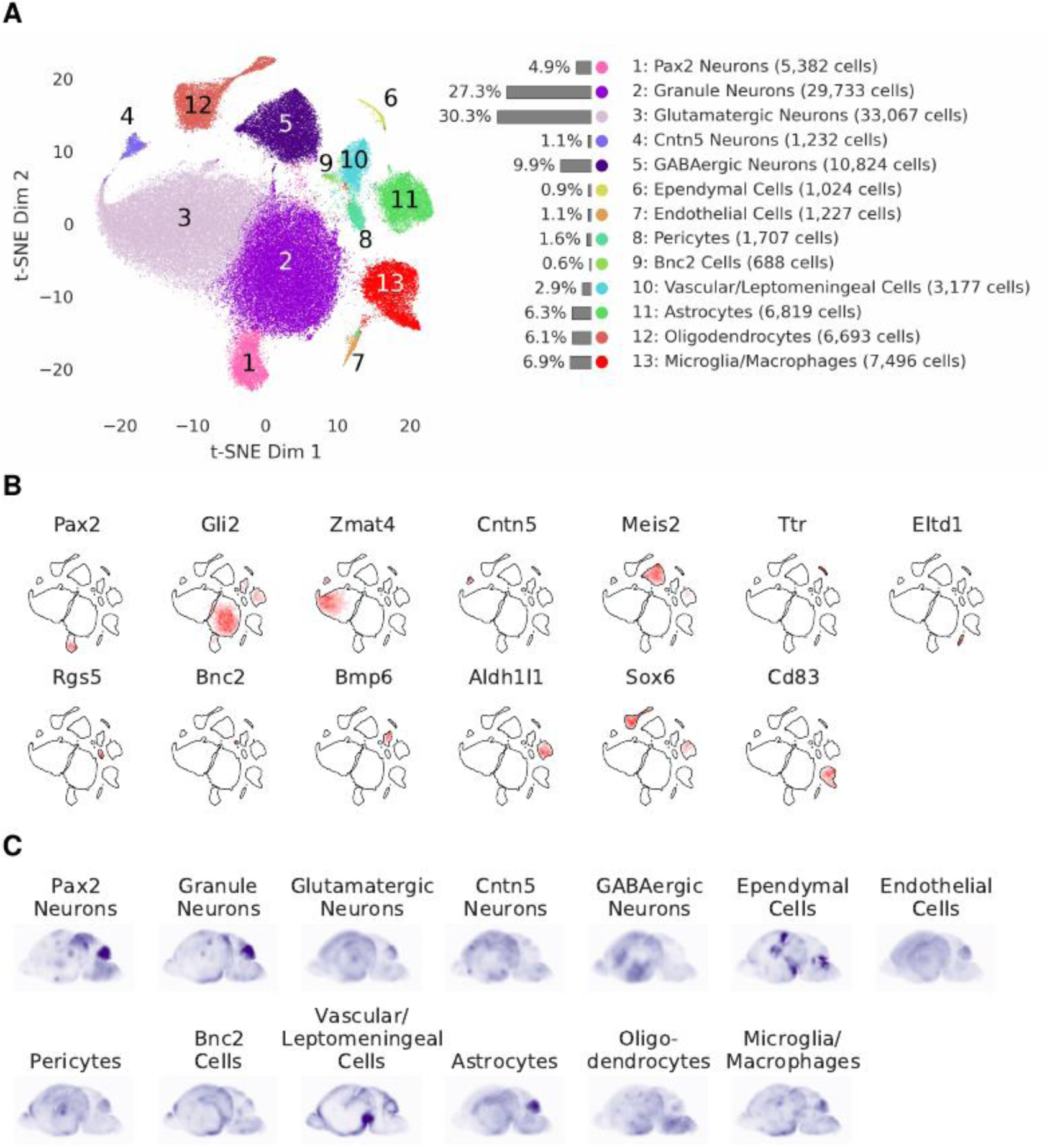
Profiling the cellular landscape of postnatal brain development with SPLiT-seq. **(A)** Profiling 109,069 cells from an entire postnatal day 5 mouse brain reveals 13 main cell types. Each cell is mapped in two dimensional space (t-SNE) and colored according to assigned cell type. **(B)** Differentially expressed genes found in each cluster correspond to putative cell type markers. **(C)** Sagittal composite ISH maps for the 13 identified cell types. For each cell type, we averaged ISH intensities across the top 10 differentially expressed genes. The displayed images are projections of several sagittal slices.

We examined *in situ* hybridization (ISH) patterns of genes differentially expressed between clusters to gain more information about the spatial origin of each cluster. We referenced the postnatal day 4 ISH data of 2,187 genes from the Allen Brain Institute’s Developing Mouse Brain Atlas (22). Composite ISH maps were generated by averaging across the 10 most highly enriched genes from each of our 13 clusters (Fig. 3C). Marker genes for granule neurons, Pax2-expressing neurons, and astrocytes were enriched in the cerebellum; those for vascular and leptomeningeal cells were enriched in the blood-brain-barrier; and those for ependymal cells were enriched around ventricular and subdural surfaces.

We then asked whether we could identify further substructure within each of the 13 main clusters of cells. Previous work has observed that t-SNE can order cells in 2D space according to stages of differentiation (10). Moving through t-SNE space along the path of differentiation can then be viewed as moving through “pseudotime” (23). When we examined the oligodendrocyte cluster, we observed a clear progression from oligodendrocyte precursor cells to newly formed oligodendrocytes, then finally to myelinating oligodendrocytes (Fig. 4A). We then characterized the temporal patterns of oligodendrocyte differentiation by measuring gene expression through pseudotime. The observed progression was similar to that reported by Marques *et al.* (10), with the exception that we did not see expression of any mature oligodendrocyte markers. This difference suggests that at five days after birth, oligodendrocyte cells have not reached maturity despite expressing myelinating markers such as *Mbp*.

**Fig. 4.**
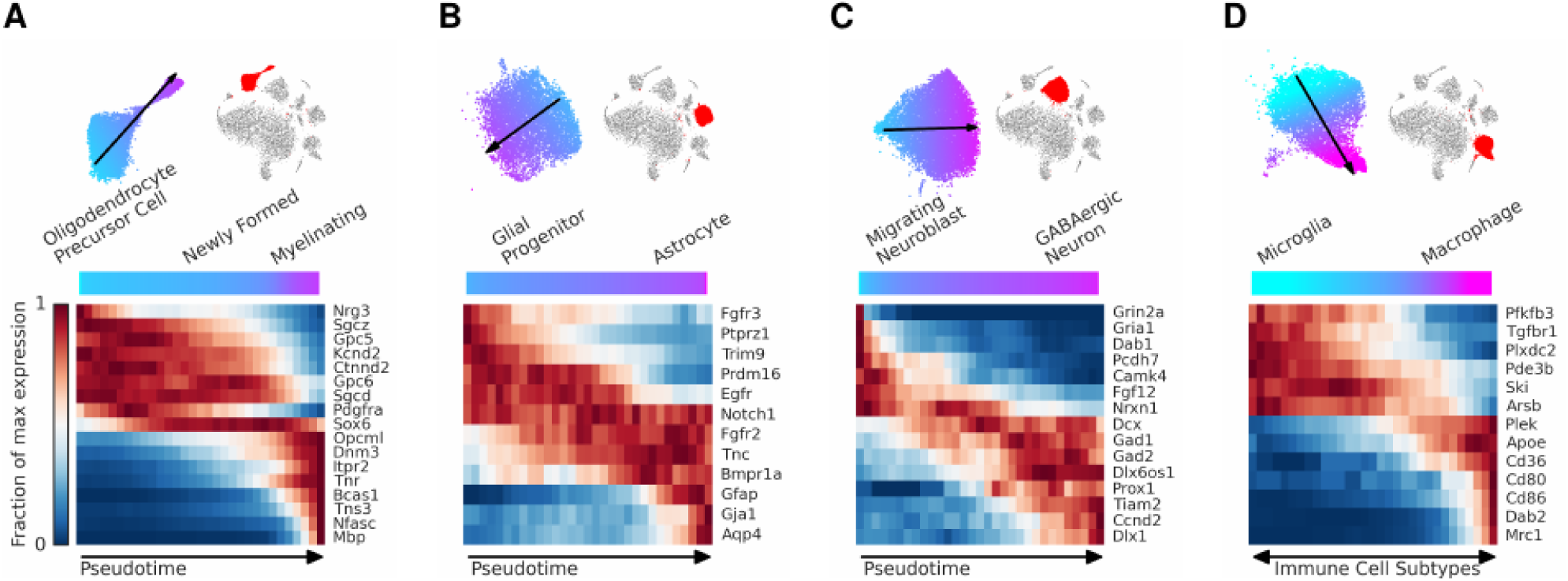
Intracluster structure reflects patterns of differentiation and subtypes present in a postnatal day 5 mouse brain. **(A)** Cells in the oligodendrocyte cluster map into t-SNE space according to their stage of differentiation. By tracking gene expression through “pseudotime” (cyan to purple), we learn the dynamics of genes during the transition from oligodendrocyte precursor cell to newly formed oligodendrocyte and then to myelinating oligodendrocytes. **(B)** Progression of glial progenitor cells differentiating into mature astrocyte cells. **(C)** Migrating neuroblasts are transitioning to GABAergic neurons. **(D)** Macrophage and microglial cells were found to populate different sides of the cluster.

Within the astrocyte cluster, cells also segregated in t-SNE coordinates according to stages of differentiation (Fig. 4B). On one end of the cluster, cells expressed glial progenitor markers such as *Ptprz1*, *Fgfr3*, and *Trim9* (24, 25), while cells on the other end expressed mature astrocyte markers such as *Gfap*, *Gja1*, and *Aqp4* (26). In the center of the cluster, cells expressed key genes in astroglial fate determination such as *Notch1* and *Fgfr2* (27, 28). There were approximately equal numbers of glial progenitors and maturing astrocytes.

A third instance of differentiation was observed in the GABAergic neuronal cluster as cells transitioned from migratory neuroblasts to mature inhibitory neurons (Fig. 4C). On the left side of the cluster, cells expressed high levels of known migratory neuroblast markers such as *Gria1*, *Camk4*, and *Fgf12* (29). Moving from left to right, expression of migratory neuroblast markers decreased as expression of known GABAergic neuronal markers such as *Dlx1*, *Gad1*, and *Gad2* (30) started to increase.

We also found two distinct populations of microglia and macrophages within one cluster (Fig 4D). Cells expressing microglia markers such as *Tgfbr1* and *Ski* (31) segregated to one side of the cluster, while cells expressing macrophage markers such as *Mrc1*, *Cd80*, and *Cd86* (32) segregated on the other side. By examining expression of these markers, we found microglial cells constituted approximately two thirds of the cluster while macrophages accounted for the remaining third.

In this work, we profiled hundreds of thousands of cells, but SPLiT-seq can be readily adapted to process many more cells. We can introduce more barcode combinations by splitting the reverse transcription into separate reactions with barcoded primers. If we were to perform this step and the three subsequent split-ligate-pool rounds in 384 well plates, we would generate nearly 22 billion barcode combinations. Additional sublibrary indexing would allow for hundreds of billions of barcode combinations, paving the way to process billions of cells in a single experiment.

Although we anticipate that the technical performance of SPLiT-seq will improve with further optimization, the current average number of reads per cell remains low compared to other scRNA-seq methods. This feature more dramatically affects cell type resolution in cases in which there are only a few, lowly expressed genes characterizing cellular diversity (*e.g*. in some neuronal subpopulations). However, for many applications, including the one presented here, there is still very valuable information to be gained. In addition, this limitation can be offset by preparing libraries from more cells (33).

The underlying methodology described here can easily be extended to new applications. SPLiT-seq’s compatibility with formaldehyde fixed cells suggests it may be able to process single cells or nuclei from dissociated formalin fixed, paraffin embedded tissue. While we profiled mammalian cells in this work, SPLiT-seq should be amenable to single-cell profiling of microbial communities as well. We also envision that SPLiT-seq can be modified to profile protein expression in single cells using multiplexed antibodies. Each antibody would be covalently attached to a DNA linker (34) that could then be barcoded with combinatorial cellular indexing. The DNA linker would also contain an antibody-specific sequence to recover the identity of the antibody after sequencing.

For scRNA-seq to become a standard tool in molecular biology labs, the barriers to entry must be reduced. Here we have presented a low-cost method to perform scRNA-seq on hundreds of thousands of cells, using only standard lab equipment. Our hope is that this increased accessibility will encourage the widespread adoption of scRNA-seq across many research areas.

## Acknowledgments

This work was supported by NIH R01CA207029 and NSF CCF-1317653 to Georg Seelig and NIH R01NS064404 and R21NS086500 to Suzie H. Pun.

